# Subspace clustering identifies transcriptome constraints determining cell type identity

**DOI:** 10.64898/2025.12.04.692451

**Authors:** Angela Huang, Junhyong Kim

## Abstract

With the advent of single-cell RNA-sequencing, researchers now have the ability to define cell types from large amounts of transcriptome information. Currently, most clustering algorithms measure cell-to-cell similarities using distance metrics based on the assumption that each cluster is comprised of “nearby” neighbors. In effect, clusters are a collection of similar cells in the embedded metric. Here, we propose that biological clusters should be comprised of sets of cells that satisfy a set of stochiometric constraints, whose intersections define a cell type. We propose to model each cell population with a single affine subspace, where all cells of the same type share a common set of linear constraints. We present an algorithm that leverages this subspace structure and learns a cell-to-cell affinity matrix based on notions of subspace similarity. We simulate scRNA-seq data according to the subspace model and benchmark our algorithm against pre-existing methods. We further benchmark our algorithm on a *C. elegans* dataset and show recovery of information on both cell type and developmental time. Lastly, we find the subspaces that our algorithm recovers allow us to find biologically significant genes involved in an organism’s development.

## I. Introduction

With scRNA-seq, researchers now have the ability to define and categorize cell types from large amounts of transcriptome information. Over the years, many clustering algorithms have been developed to automate this task. Motivation for such work can be seen in various cell atlas projects (e.g. the Mouse Cell Atlas [1][2] and Human Cell Atlas [3]), which aim to depict the cell types present in an organism across its various stages of development. Not only does this give researchers a deeper understanding of the biological complexity of the organism, but it also serves as a reference for disease studies.

Given their potential impact, the single cell literature is rich with different unsupervised clustering techniques [4][5][6]. Most popularly, iterative clustering methods such as k-means/Lloyd’s algorithm [7][8][9][10], hierarchical clustering-based algorithms [10][11] [12][13], and community-detection based algorithms [14] [15] have been widely employed. Spectral clustering and low-rank methods have also been proposed [16] [17]. Each clustering category proposes its own model of the data. For example, distance to cluster centroid methods (such as *k*-means) assume roughly isotropic clusters with centroids. The most common hierarchical clustering methods (Wards and “complete” linkage) also assume compact clusters around cluster centroids. Density based methods typically do not assume clusters of a particular shape of size but assume that all clusters are equally dense or in biological terms, all cell types are equally homogenous. Graph based methods are an extension of density-based methods. They usually exploit a notion of “modularity” or sparsity, and cell types of different sizes, densities, and shapes can be identified. More recently, deep learning methods involving autoencoders that optimize reconstruction accuracy and clustering accuracy have additionally been gaining popularity [18][19][20][21].

Despite having different model assumptions, most of the clustering frameworks mentioned above share a common theme: After the optional step of performing dimension reduction, the methods measure cell-to-cell similarity using distance metrics defined on the ambient feature space. Therefore, they often work best when cells of the same type (cluster) are relatively close in the ambient space, and cells originating from different tissues/types are relatively far apart. This poses a general constraint that the clusters be both compact and far enough apart that the algorithms can recover them.

In practice, this may not always be the case. Consider developmental data, for example. The underlying clusters may not be compact; the cells may be well spread out across developmental trajectories (Fig.1c). The clusters may also not be very far apart; the developmental trajectories may intersect (Fig.1b). More importantly, computing distances in the ambient space implicitly assumes that the entire measurement space, i.e., the entire transcriptome, is relevant to every cell type. This ignores the fact that different cell types might have different dimensionalities due to different pathways being differentially on or off. In fact, we propose that a cell type, arising out of its molecular microstate, is not determined so much by exact combination of molecular components. Rather, we propose that cells “occupy” a permissible ensemble of molecular states that satisfy physiological and biochemical reaction constraints. That is, as long as the cell satisfies reaction constraints for a given cell function, it will manifest phenotypically as a particular cell type. Within the mRNA state space, this predicts that a cell type is expected to lie on a lower dimensional subset of possible transcriptomes defined by biochemical flux balance of proteins and functional RNA [22][23][24]. At equilibrium flux balance relationships will create stochiometric relationships, which in molecular state space will define geometric subsets generated by low order polynomials (i.e., algebraic varieties [25]), which represent molecular interaction orders.

**Fig. 1.**
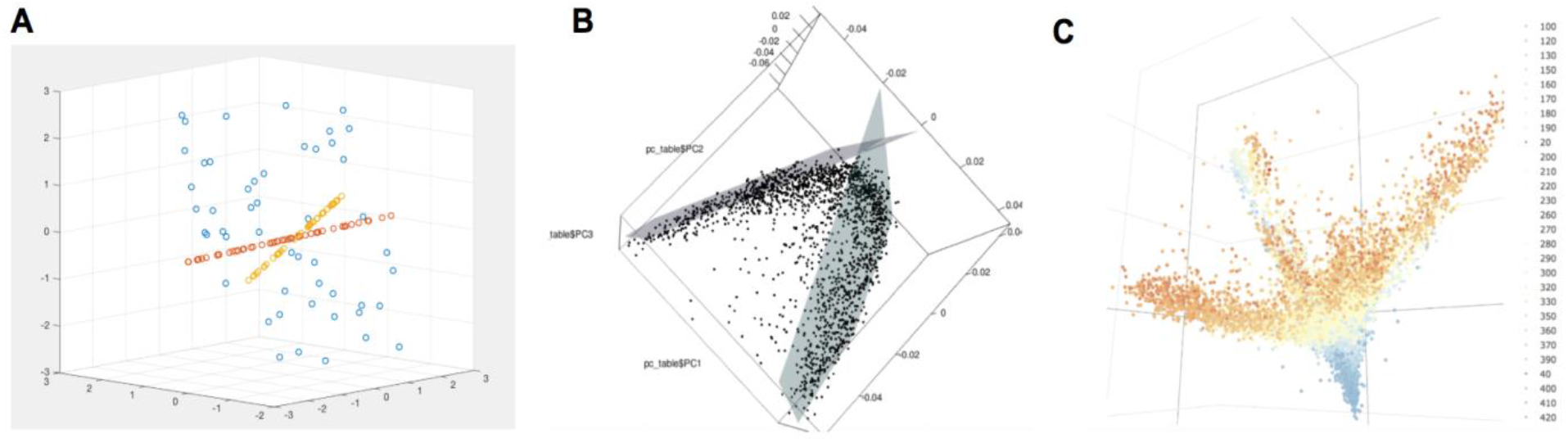
Visualizing the underlying subspace structure in the simulated and real datasets. (A) PCA plot of the low-dimensional simulated dataset; (B) PCA plot of the body wall muscle cells of the *C. elegans* dataset, demonstrating early cells residing in one subspace and later cells residing in another; (C) PCA plot of the full *C. elegans* dataset, colored by time where blue points represent cells at an earlier time-point and orange points represent cells at a later time-point

**Fig. 2.**
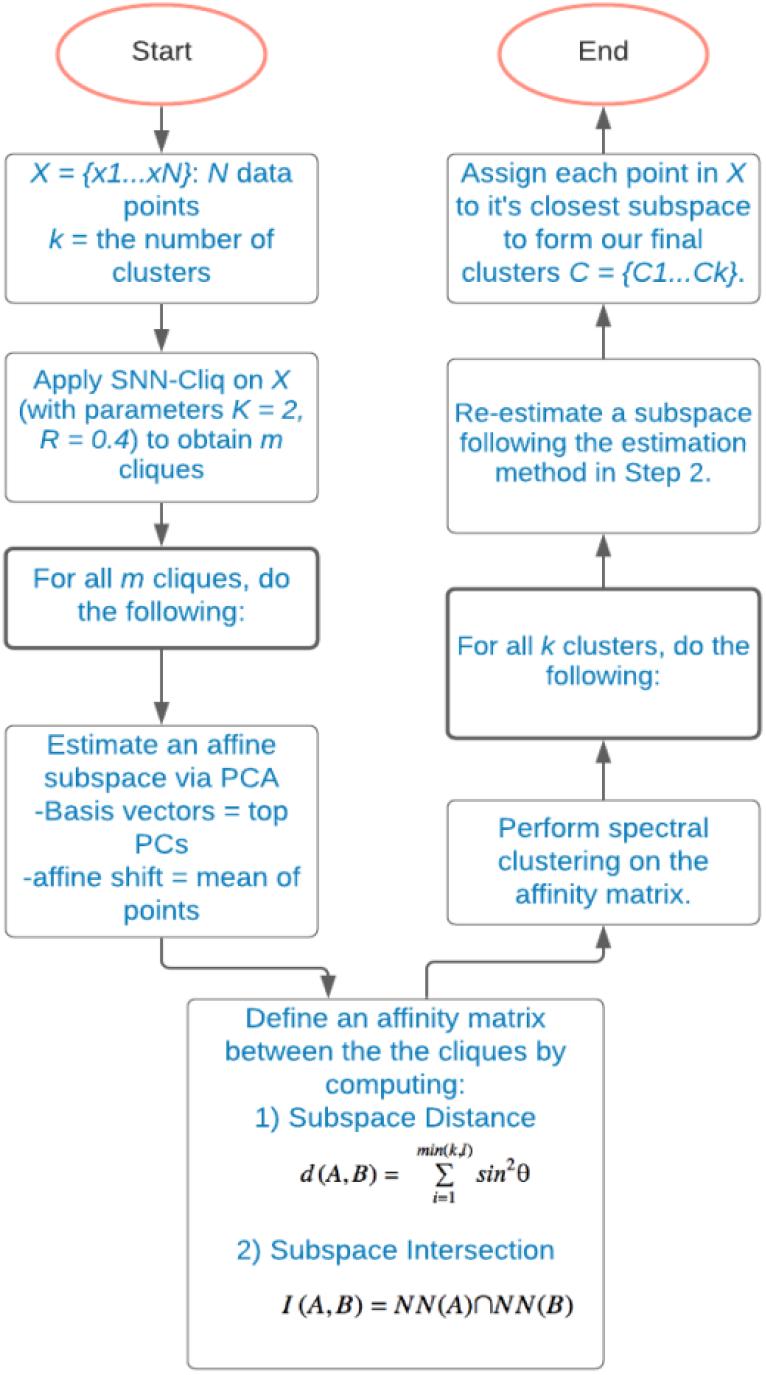
A flow chart describing the subspace clustering algorithm

Given the dataset noise and computational complexity when it comes to identifying polynomial subsets, we instead consider clustering cell types as individual subspaces of the ambient space. We can think of such subspaces as convex bounds on the complex manifolds that characterize a cell type’s molecular physiology. With this idea, we choose to group cells based on similarities along linear combinations of a gene expression subspace, rather than similarities in pairwise gene distances. We assume each cell type lies in a separate affine subspace. Then, recovering the various cell types present a dataset reduces to the problem of subspace clustering: Given a dataset that lies near a union of affine subspaces, simultaneously cluster the data into multiple subspaces while finding a low-dimensional subspace that fits each group of cells. In general, finding *k* distinct subspaces so as to minimize the distances between the given points and their respective nearest subspaces is NP-hard [26]. To address this, we develop a bottom-up heuristic approach to solve our task. We apply our subspace clustering algorithm to simulated datasets as well as an in-house *C. elegans* dataset [27]. We also benchmark the algorithm against other popular algorithms in the field. We found this algorithm has the ability to recover cell types present in an organism, cell subpopulations, and differentiate cells according to their developmental time. Lastly, we found the subspaces our algorithm recovers allow us to find biologically significant genes involved in an organism’s development.

## II. Method

We model the dataset *X* ∈ ℝ^*D*^ as a union of affine subspaces, where each subspace corresponds to a cell type. Let us denote these subspaces as *S*_*i*_ where *i* = 1 … *K*.

The goal of our subspace clustering algorithm is four-fold:

1. *Find the number of subspaces: K*
2. *their dimensions: d*_*i*_
3. *a set of basis vectors and an affine shift for each subspace: B*_*i*_ *and a*_*i*_
4. *the segmentation of cells to their nearest subspace: C*_*i*_.

### 1: Initialize the search space

As a first step in our bottom-up approach, we aim to find small groups of homogenous cells. To establish vocabulary, we call a group of cells *homogenous* if they all belong to the same cell type. We find dense regions of cells in the dataset, by leveraging Xu et al.’s Shared Nearest Neighbor Cliq algorithm which utilizes shared nearest neighbor metrics to build a connectivity graph and then partitions the graph through a quasi-clique finding approach [28]. Setting the algorithm’s parameters to their most stringent settings for the dataset helps ensure that the dense regions we recover are homogenous.

### 2: Estimate an affine subspace for each clique

Next, we perform PCA on the points in each clique to estimate an affine subspace for each clique from step 1. We assign the principal components as the basis vectors of the subspace and the mean of the points as the affine shift. To determine the subspace dimension, we generate subspaces of varying dimension by incrementally increasing the number of top principal components obtained from the original PCA. Among these subspaces of varying dimension, we find the one with dimension *d* such that the projection of the original points onto this subspace yields a reconstruction accuracy of 95%, as computed using the eigenvalues of the covariance matrix of *X*: *λ*_1_ + ⋯ + *λ*_*d*_ / *λ*_1_ + ⋯ + *λ*_*n*_.

### 3: Define an affinity matrix between the estimated subspaces

We next utilize two metrics of subspace similarity to identify ensembles of points in same subspace by building an affinity matrix using these metrics.

#### a) Subspace Distance

Ye-LHL[29] showed that any valid distance function *d*(*A, B*) on subspaces *A* of dimension *k* and *B* of dimension *l* must be a function of only their principal angles *θ*_1_ … *θ*_*min* (*k,l*)_. Therefore, we use the *Chordal* distance on principal angles as our first similarity metric:

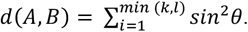

#### b) Subspace Intersection

This metric measures the number of overlapping cells between subspaces. Let *NN*(*A*) be the closest *x* points to an arbitrary subspace *A* and *NN*(*B*) be the closest *y* points to an arbitrary subspace *B* using the cumulative sum method on distances [30]. Then compute:

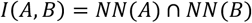

We construct an *m* × *m* affinity matrix by computing pairwise affine subspace distances and pairwise affine subspace intersections and summing the values together.

### 4: Perform spectral clustering on the affinity matrix

Then, we perform spectral clustering to group the *m* cliques into *K* clusters of cliques.

### 5: Re-estimate a subspace for each cluster following the estimation method in Step 2

### 6: Assign each point *x* ∈ *X* **to its nearest subspace using the Manhattan distance to form our final clusters** *C*_1…_*C*_*k*_

## III. Results

To validate the performance of our algorithm, we test our algorithm on simulated datasets and then demonstrate an application using in-house *C. elegans* dataset. First, we curate a simulated dataset by generating points that lie on low-dimensional affine subspaces to show that the subspace clustering algorithm works on our target geometric model. Next, we simulate a more complex dataset by using estimated subspaces parameterized from a *C. elegans* dataset using the original papers’ cell type and developmental stage classifications [27].

For our simulation, for each given subspace *S*_*i*_ with dimension *d*_*i*_, we generate points in ℝ^*D*^ following this formula:

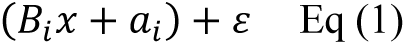

- 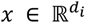 is a low dimensional vector representation of the point
- 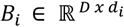 is the set of basis vectors of the subspace
- *a*_*i*_ ∈ ℝ^*D*^ is the affine shift of the subspace
- *ε* ∈ ℝ^*D*^ is a noise vector describing the distance of the point from the subspace

### A. Synthetic Dataset 1: Low-Dimensional Affine Subspaces

Data points were uniformly generated on two random affine lines (subspaces of dimension 1), and a random affine plane (subspace of dimension 2). The clusters are of equal sizes and contain 50 data points each. In our benchmarking analysis, we consider a variety of single-cell algorithms that fit different model assumptions. Standard *k*-means and pcaReduce are *k*-means based methods, Louvain and Leiden (as implemented in Seurat) as well as SNN-Cliq are graph-based methods, CIDR employs hierarchical clustering, pcaReduce combines *k*-means and hierarchical clustering, and SIMLR and SinNLRR are spectral methods. Due to numerical errors when running single-cell methods designed for high-dimensional datasets on very low-dimensional datasets, we test a subset of our benchmarking methods here. We measured the performance of the clustering with three evaluation measures: Adjusted Random Index (ARI), Normalized Mutual Information (NMI), and Purity scores [31][32][33]. As can be seen in Fig. 3 and 4, our bottom-up subspace clustering heuristic algorithm accurately identified subspace clusters, even when the points from different clusters closely overlapped in Euclidean space (Fig.3). Other clustering methods including an explicit low-rank clustering method SinNLRR did not perform as well (Fig.4a).

**Fig. 3.**
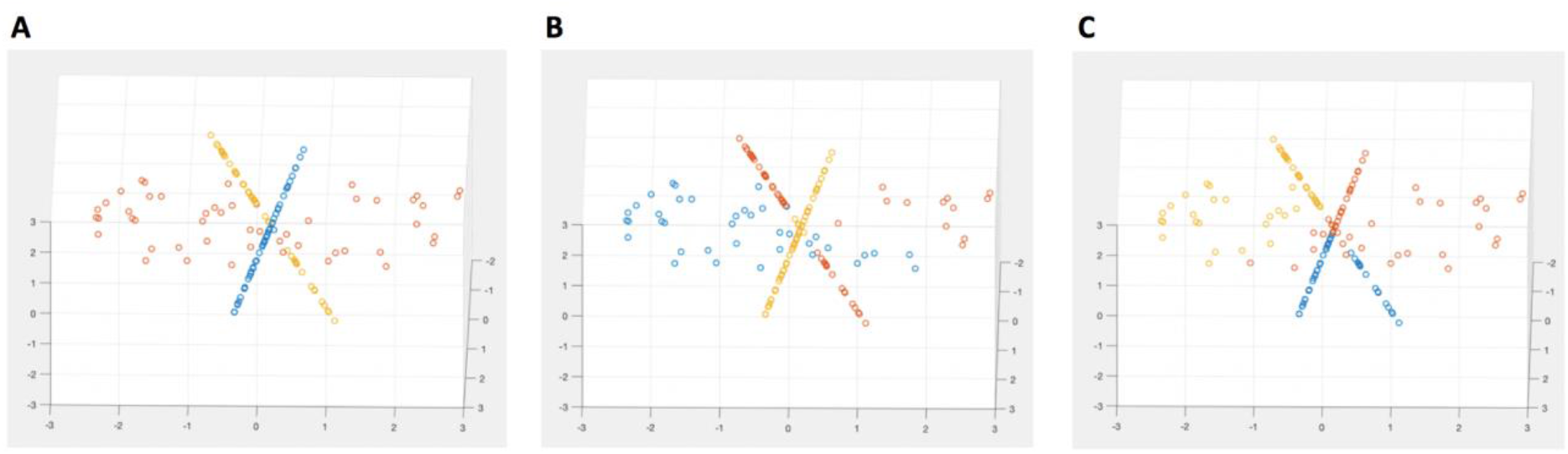
Visualizing the clustering results of various algorithms. (A) Subspace clustering method correctly recovers the two lines and planes; (B) SIMLR is able to piece together the lines but fails to recover the sparser plane; (C) SinNLRR does not fully recover any of the low-dimensional subspace structure but clusters the points in their local regions

**Fig. 4.**
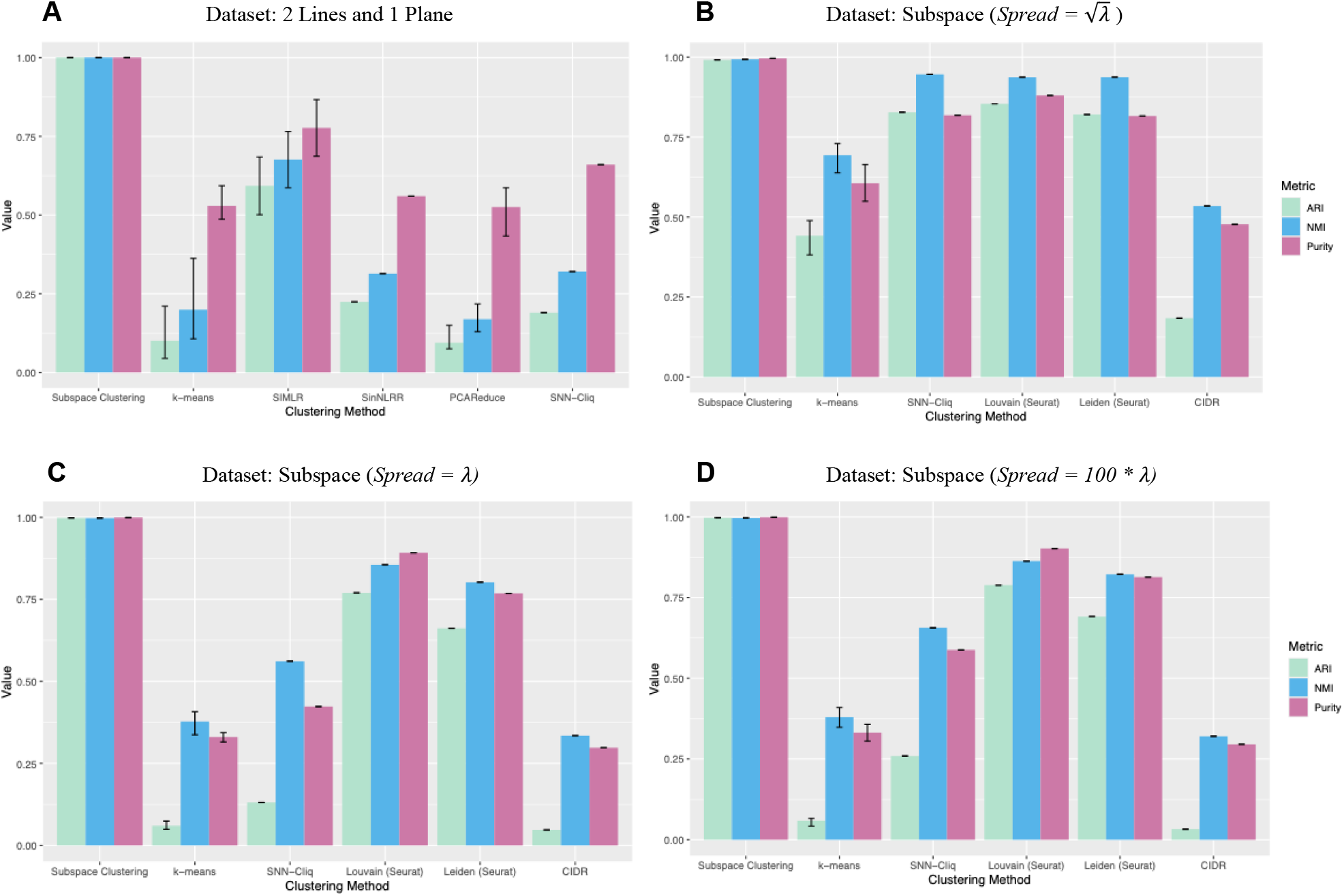
(A) Clustering results on synthetic dataset 1 containing low dimensional subspaces. Each algorithm was run 20 times and the means are shown. The error bars reflect the minimum and maximum scores over the 20 runs. (B) - (D) Clustering results on synthetic datasets 2-4 containing 11 cell types. Each dataset contains 2200 cells with 200 cells per cell type. The data are modeled as belonging to affine subspaces. The vector *x* varies across each dataset. The error bars reflect the minimum and maximum scores over 10 runs.

### B. Synthetic Datasets 2-4: Clustering 11 Cell Types modeled as Subspaces

To create a more data-driven simulation, we analyzed the single cell empirical dataset in [27] to estimate subspaces for each cell type, using the annotations given in the paper, and then simulate points onto these subspaces using the Eq (1) model. We chose 11 major cell types: Arcade Cell, Body Wall Muscle, Ciliated Neuron, Non-Ciliated Neuron, Coelomocyte, Glia, Hypodermis, Intestinal and Rectal Muscle, Intestine, Pharyngeal Gland, and Seam Cell. We choose these particular cell types because they are biologically diverse, contain enough cells for the algorithms to recover them, and include a variety of subspace dimensions.

We generated three datasets to model increasing spread by varying the scale of the low-dimensional vector *x*. We first estimate the spread along each subspace direction from the real dataset by observing the eigenvalues *λ* corresponding to the estimated subspace basis vectors. For each basis vector, we draw a random value from a normal distribution modeled with a mean of zero and standard deviation equal to the square root of the corresponding eigenvalue or 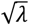 as this represents the standard deviation of the data along the particular eigenvector direction in the real data. We assign these values to the vector *x*. For the remaining two datasets, we then increase the spread by assigning *λ* and then 100* *λ* as the standard deviations of the normal distribution respectively to determine the impact of increasing geometric spread and therefore sparsity on clustering performance.

Our results show that even under this complex setting with 11 different cell types, our algorithm was able to almost perfectly recover all 11 simulated cell types. As the spread increased, the performance of the subspace clustering algorithm remained consistent, while the performance of most other methods decreased. *K*-means fails to recover the 11 cell types in all datasets due to its inherent model assumption that the clusters are roughly isotropic clusters with centroids. SNN-Cliq works relatively well on the first dataset but its performance quickly deteriorates with increasing spread and sparsity. The Louvain and Leiden algorithm as implemented in Seurat perform similarly across the three datasets but in all instances are unable to recover all 11 cell types. (Fig. 4b-4d)

### C. Synthetic Datasets 5 and 6: Clustering 11 Cell Types modeled as Mixtures of Gaussians Subspaces

We further generated two datasets to represent a mixture of multivariate gaussians rather than affine subspaces Again, we utilize the empirical dataset in [27] to guide our simulations and the same 11 major cell types were chosen. In our first gaussian mixture dataset, we assign the estimated affine shifts of the subspaces as the means of our gaussian clusters and utilize the covariance matrix calculated from the real data for each cell type. In the second dataset, we again assign the estimated affine shifts of the subspace as the means of our gaussian mixture models clusters but assign the identity matrix as the covariance matrices to eliminate the underlying unknown relationships between genes that could largely impact the clustering results.

**Fig. 5.**
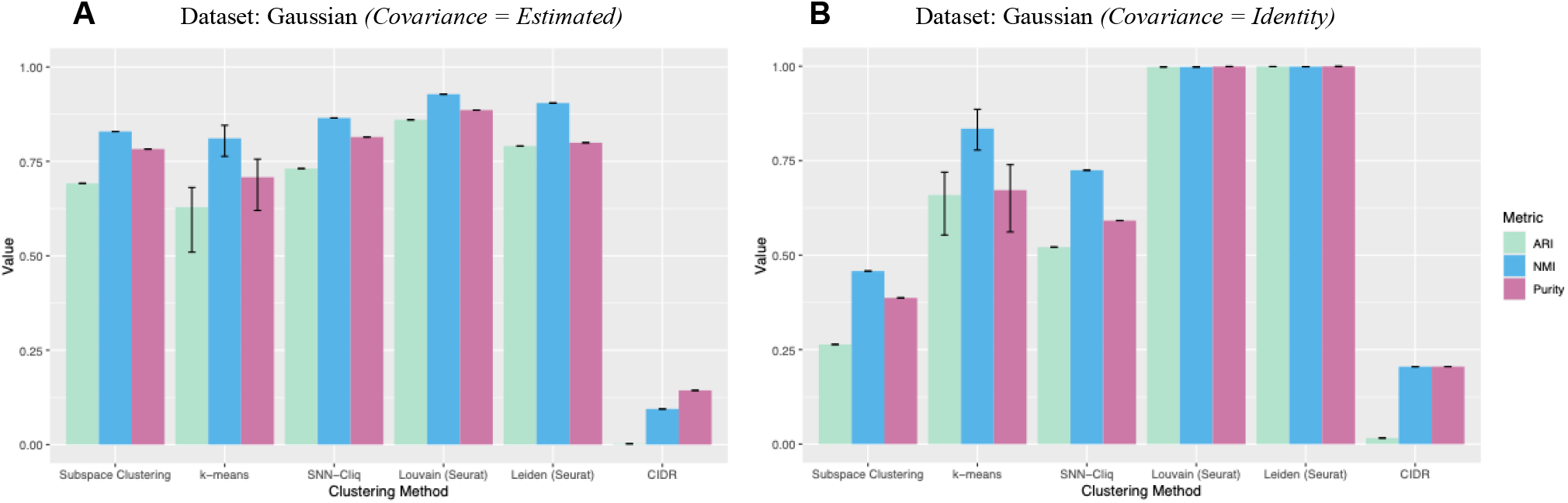
Clustering results on the simulated datasets containing 11 cell types. The data are modeled as a mixture of gaussians. Each dataset contains 2200 cells with 200 cells per cell type. The error bars reflect the minimum and maximum scores over 10 runs. (A) The covariance matrices are calculated from the real data. (B) The covariance matrices are identity matrices to eliminate the underlying unknown relationships between genes that could largely impact the clustering results.

Overall, popular methods such as k-means, Louvain, and Leiden performed better on the mixture of gaussians datasets than the subspace datasets, especially when the identity matrix was introduced as the covariance matrix. This confirms the idea that current methods generally work well on simpler, well-studied models such as the gaussian mixture model but have difficulty in more complex cases such as when clusters conform to a mixture of affine subspaces. Furthermore, when introducing the identity matrix as the covariance matrix, this effectively increases the dataset noise for the subspace clustering algorithm and its performance decreases as expected. At the same time, replacing the estimated covariance matrix with the identity matrix essentially reduces the dependencies between genes and we see an overall improvement in performance for methods including *k*-means, Louvain, Leiden, and CIDR. The difference in performance between the two Gaussian datasets suggest that not all genes are equally important and that there are underlying relationships between genes of varying degrees that influence the clustering results.

### D. Performance on C. elegans Dataset

Here, we tested the algorithm’s ability to cluster the original 34 major cell types annotated in the *C. elegans* dataset, and we benchmark our method against other single-cell clustering algorithms. As part of our preprocessing steps, house-keeping genes and cell cycle genes were removed. The dataset was size-factor normalized, background corrected, and log transformed. Intermediate cells between developmental timepoints 400-500 in the original dataset were removed. To reduce computational complexity, the top 6752 variably expressed genes (VEGs) were selected for analysis.

Overall, the method that performed the best across all metrics on the real dataset was Louvain as implemented by Seurat. The subspace algorithm came close to and then matched Seurat when considering the NMI and Purity scores respectively. One reason for this is the so-called ‘‘double-dipping’’ [34] or ‘‘selection-bias’’ problem. During the data annotation stage, a rough clustering of the *C. elegans* dataset was first derived using popular algorithms including Seurat and afterwards each cell was manually annotated using domain knowledge. In effect, the dataset labels may be unfairly optimized for the specific criteria and assumptions of the algorithm it was trained on, leading to possible inflated performance results for these methods during the benchmarking stage. When looking at the algorithm’s behavior, the subspace clustering algorithm tends to break up clusters into smaller, but still homogenous clusters, so it tends to perform on par with the highest performing methods when evaluated using the NMI and Purity metrics but poorly when evaluated with ARI. In fact, due to developmental time present in the dataset where we observe that early cells often occupy one subspace and later cells shift into another subspace across developmental time, the subspace clustering performance increases to ARI of 0.33, NMI of 0.76, and Purity score of 0.87. In general, we would expect the NMI and Purity scores for a specific algorithm to be similar to each other because they compute the accuracy of the final clustering similarly, but NMI adjusts for cluster size distribution.

**Fig. 6.**
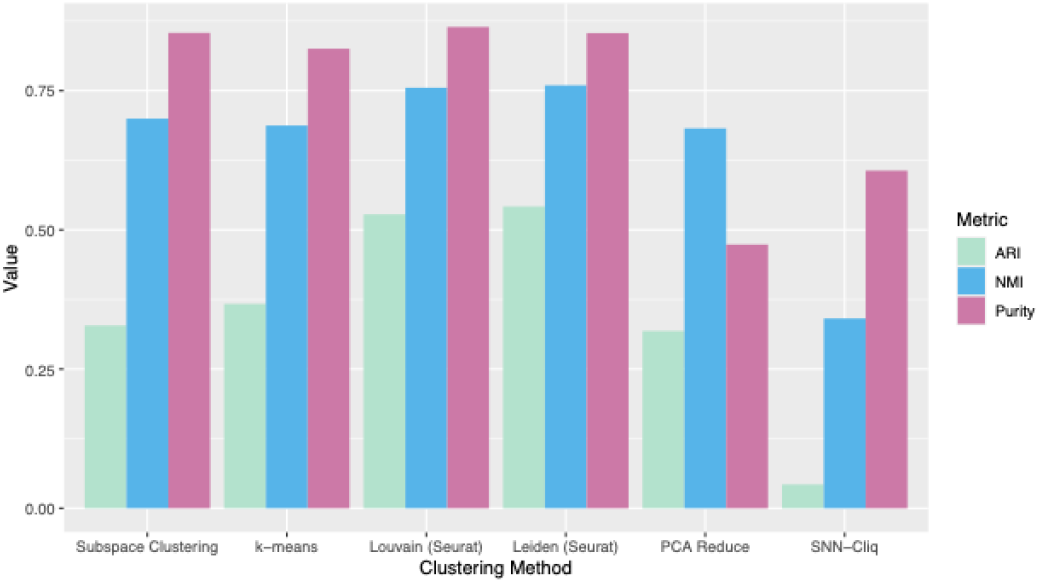
Clustering results on the *C. elegans* dataset with 34 cell types.

### E. Application to Learning Separating Hyperplanes

One utility of the subspace clustering method is that it allows us to identify biological processes where cells show abrupt shifts in subspace relationships due to large-scale changes in pathway usage as might happen within a cell type during different stages of development (Fig.1b). Here, we ran the subspace clustering algorithm on three major cell types in the *C. elegans* dataset and set the parameter for the number of clusters to two. Early cells were defined to be within the time range 300-400, and late cells were defined to be in the time range: >500. We then calculated how well the algorithm performed on clustering the early versus late cells. When clustering cells in the Body Wall Muscle, Hypodermis, and Ciliated Neuron into early and late hyperplanes, the algorithm consistently achieved ARI, NMI, and Purity scores above 0.998. For the Arcade Cell, the subspace clustering algorithm achieved an ARI of 0.901, NMI of 0.8476, and Purity score of 0.918. For clustering the Intestine, the algorithm achieved an ARI of 0.954, NMI of 0.892, and Purity score of 0.968. Using our subspace method, we were able to accurately identify early and late-stage differences in cells of the same type

### F. Finding the Major Genes Constrained by the Orthogonal Complements of Subspaces

As noted in the introduction, our rationale for subspace clustering is that cell types occupy subspaces as a result of satisfying stochiometric constraints. Here, we attempt to find the major genes defining the subspace of a cell type by exploring the notion of orthogonal complements (Fig.7). We propose that a linear combination of genes are significant in constraining cells to lie in a certain region of the high dimensional space as a result of various biochemical constraints. These constraints are expected form invariants, that is directions of variation that are constants or zeros within the identified cell type subspaces. We propose that these constraint sets will belong to the orthogonal complement of the cell type subspaces.

**Fig. 7.**
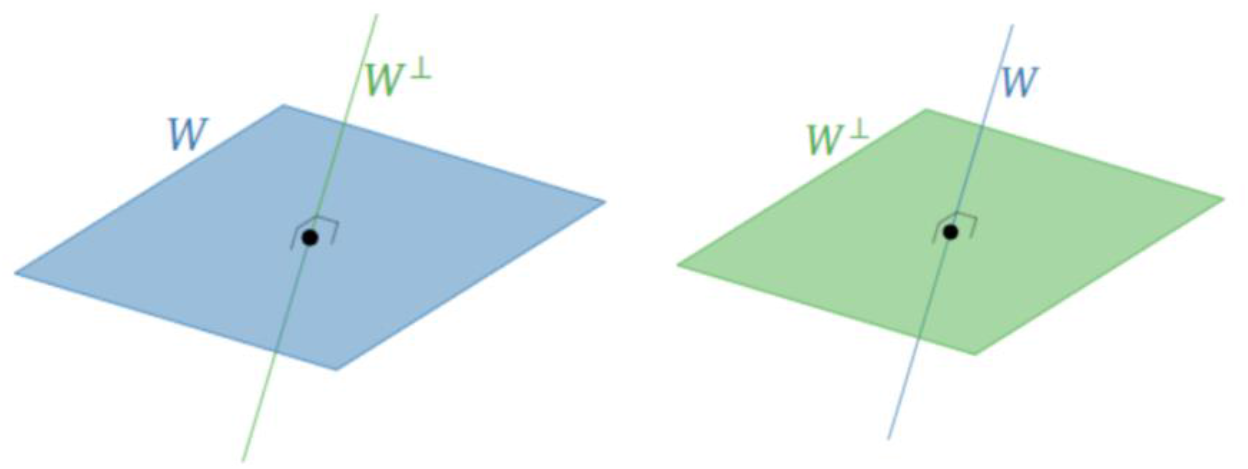
The orthogonal complement of a plane *W* in ℝ^3^ is the perpendicular line *W*^⊥^. The linear combination of genes in ℝ^3^ that make up *W*^⊥^ constrain cells to lie in the plane *W*. The orthogonal complement of the line *W* in ℝ^3^ is the plane *W*^⊥^. The linear combination of genes in ℝ^3^ that make up make up *W*^⊥^ constrain cells to lie in the line *W*. [35]

To find the major genes in the cell type subspace that are constrained by the orthogonal complement, we run sparse principal components analysis (SPCA) on each cell type in the dataset. SPCA is a variant of PCA that aims to return sparse loadings to concentrate the basis vectors to a small number of major effect loadings. The ‘sparsepca’ library in R was used for this task. After running SPCA and finding a sparse subspace with dimension *d*, we then look at the top two principal directions of the estimated subspace for the cell type and record the genes that are present in these loadings. As an alternative approach to identifying genes involved in constraining the cell type to particular subspaces, we also take a gene-centric approach and search for the genes within a cell type with the highest expression and lowest variance. These are genes that when projected onto the cell type subspace, they tightly group together and exhibit low variance, i.e., their values are constrained. We add the criterion of high expression to select genes that are active but constrained. The general idea is that genes should display high expression values indicating they are present and important in defining a particular cell type, but the variance in the gene’s expression should be low due to stringent biochemical constraints. (Fig. 8)

**Fig. 8.**
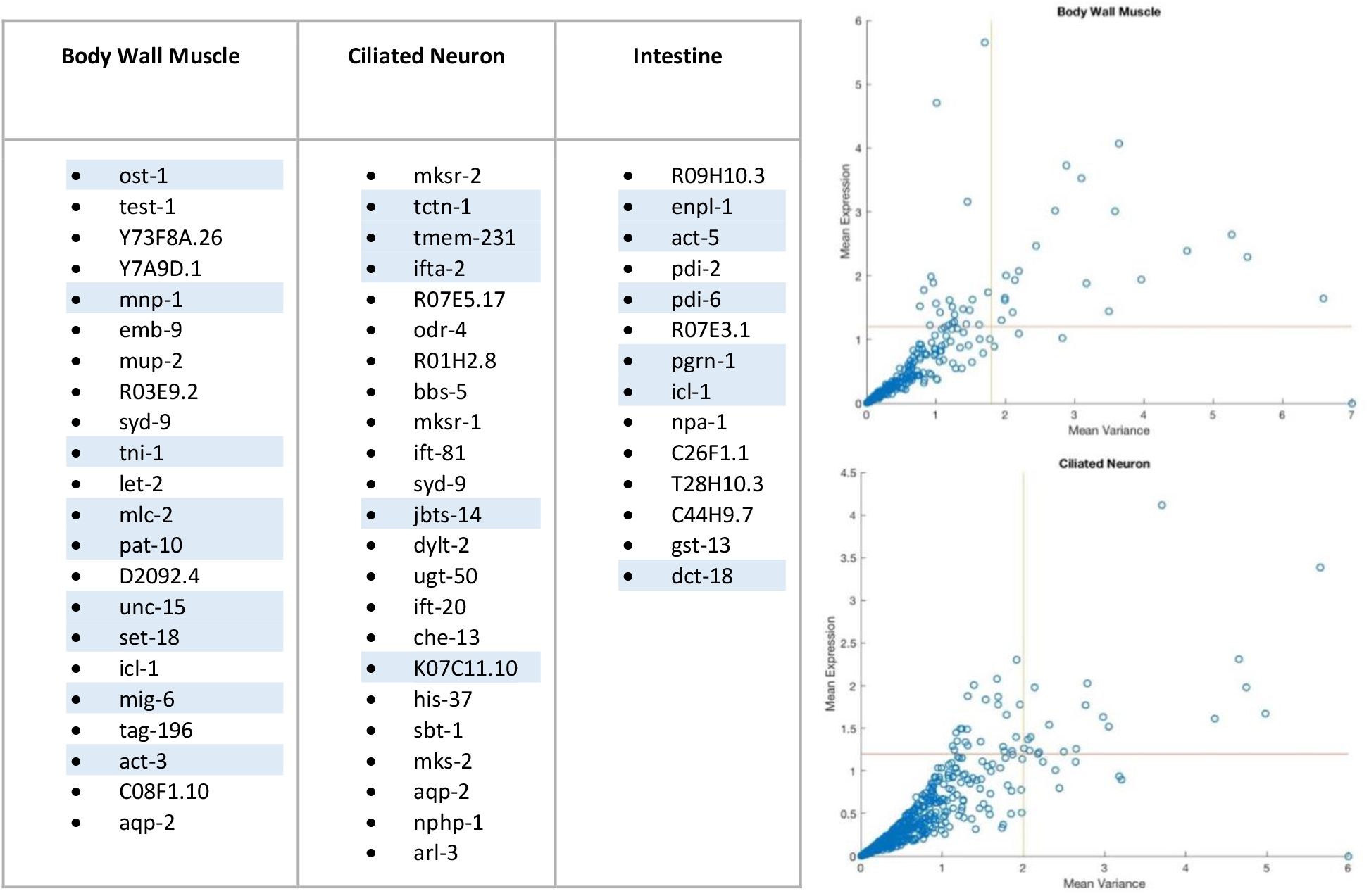
A table of the genes for each cell type with high expression, and low variance. Gene annotations for each gene were queried using WormBase. Genes that are enriched in their respective cell types are highlighted in blue. The variance vs. mean expression plots for two cell types are depicted on the right.

To compare our “cell-type defining” gene list to an existing knowledge base, we curated a list of “literature” genes by parsing WormBase’s database of gene annotations. For each cell type, we extracted all relevant genes containing annotations that state that the gene participates in the function of the particular cell type. For each of our two lists of cell type defining genes, we compared their intersection with the WormBase literature set. We further compare our gene sets with the top 20 genes obtained from Seurat’s marker genes finding method which returns the genes with the highest average log-fold change using the non-parametric Wilcoxon rank sum test. For each cell type and gene list, we computed an expected *literature gene ratio* which is the probability of randomly selecting a gene that is enriched in a particular cell type. This is calculated by taking the size of the literature list for that cell type and dividing it by the total number of genes in the dataset.

Across three major cell types, we find that we recover significantly more genes than Seurat from the literature set using our subspace-oriented gene finding methods (Fig. 9). Using a proportion test, we also find that the proportion of cell type-relevant genes we recover in our gene finding methods are significant compared to randomly selecting a set of genes (the *literature gene ratio*).

**Fig. 9.**
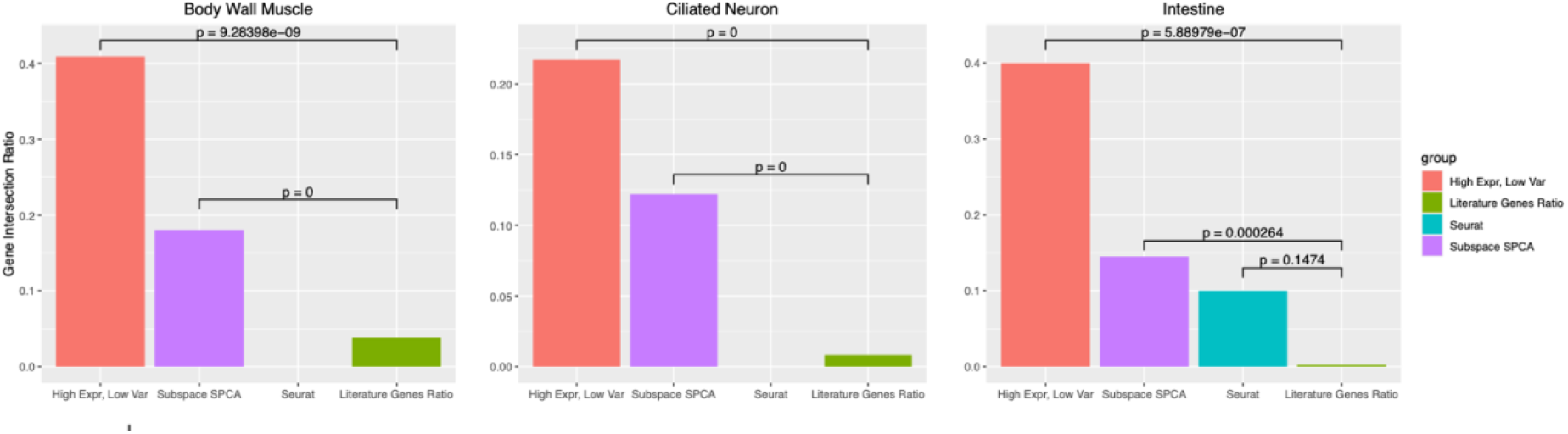
A bar chart comparing the gene intersection ratio for three major cell types. The gene intersection ratio is calculated as the number of genes found in the intersection of the literature gene list and derived gene list, divided by the size of the derived gene list. Two gene lists were derived using Sparse PCA and the gene-centric high expression, low variance orthogonal complements analysis. We compare our gene finding methods with the top 20 genes obtained from Seurat’s marker genes finding method. We then obtain a *literature gene ratio* which is the probability of randomly selecting a gene that is enriched in a particular cell type. This is calculated by taking the size of the literature list for that cell type and dividing it by the total number of genes in the dataset. We then compute the significance of the cell type-relevant genes we recover in the various gene finding methods compared to randomly selecting a set of genes (the *literature gene ratio*). These are shown by the p-values displayed in the chart.

**Fig. 10.**
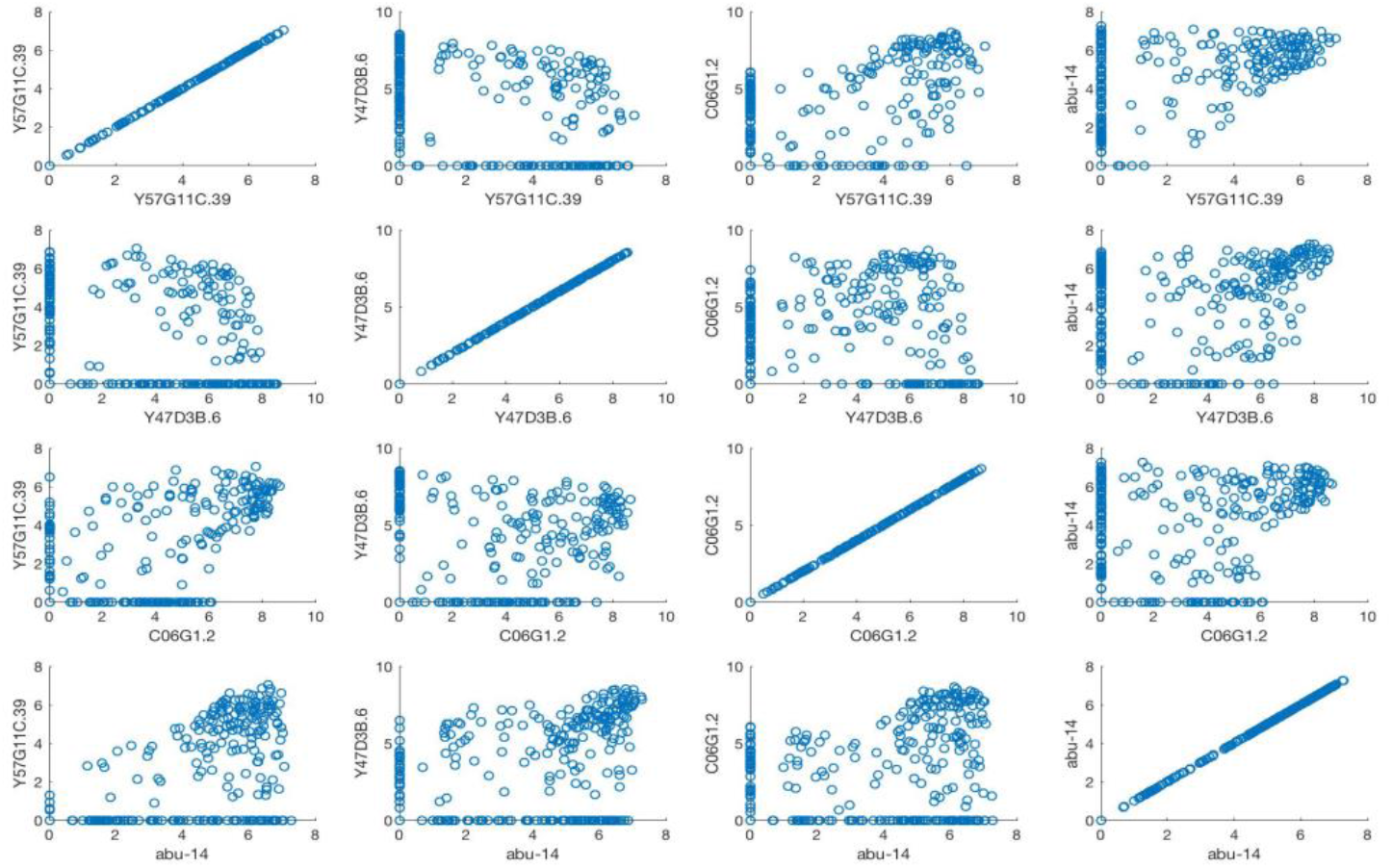
A matrix plot illustrating gene *x* gene interactions in the first orthongonal complement loading of the Arcade Cell. See the Appendix for matrix plots of other cell types.

To get a sense of how the genes interact, we further depict the gene by gene interactions in a matrix plot for all the cells within a cell type.

## IV. Discussion

Overall, we show the subspace clustering method to be a promising technique for identifying cell types in an organism under the assumption that different cell types occupy different manifolds or algebraic varieties, which are approximated by affine subspaces. It also identifies major shifts in cell states through temporal dynamics such as during development. We further show that the conceptual approach of identifying constraining set of genes, rather than simply active genes, is more powerful in inferring key genes that determine cell identity.

As noted in the introduction, the problem of identifying optimal subspace clusters is NP-hard and we developed a bottom-up heuristic method. Several factors affect the subspace identification. First, the *amount of high dimensional noise* in the data is a key determinant of performance. This problem is a fundamental aspect of all dimension identification problems and the subject of continued research [36][37]. Second, the configuration of the subspaces occupied by the cell type states determine effective cluster recovery. The *principal angles* between clusters measures how much the clusters differ in directions, which in biological terms refer to similarity of covariation of genes. As this value becomes smaller, the clustering problem becomes harder. Additionally, as the *number of overlapping dimensions* (i.e., overlapping gene pathways) between the subspaces increases, the amount of separation between clusters decreases, making it harder for algorithms to differentiate between clusters. Lastly, the location of the *affine shifts* of the subspaces, representing overall mean expression difference, also plays a role, albeit this is also key for all clustering algorithms. While these factors all affect subspace clustering performance, our simulation studies show good performance in our ability to recover true subspace clusters. Our algorithm was especially effective for the difficult cases presented in Figure 4.

What determines a cell type is both an empirical and a conceptual problem [38]. Regardless of whether cell types are determined by molecular physiological constraints, it is well established that different cell types express different numbers of genes [39] and different pathways, implying that they “live” in different dimensions. Measuring differences for objects that have different intrinsic dimensions is a difficult problem and likely to distort any standard clustering method that assume full measurement space. Furthermore, we propose that well-appreciated single cell variation arises not just from stochastic noise, but because system drift is allowed as long as biochemical constraints related to cell phenotype is maintained as previously proposed in flux-balance type of models [22]. This suggests the subspace model to be a promising model for partitioning and recovering different cell types that lie on the single-cell manifold. Future work in this direction includes considering more bounded subspaces to incorporate the biologically realistic assumption that expression range is bounded, incorporating pathway knowledge base into constraint set identification, and speeding up the algorithmic approaches to scale to millions of cells that are now available. In sum, we propose that the idea of clustering by biologically feasible geometric sets is appropriate for modeling cell phenotypes within single cell distributions.

## Acknowledgements

This work was funded by Health Research Formula Fund from the Commonwealth of Pennsylvania who had no direct role in the work. The authors have no competing interests. Source code for the algorithm can be found on Github. The raw data has been deposited in the Gene Expression Omnibus (**www.ncbi.nlm.nih.gov/geo**) under accession code GSE126954.

